# Hepatocyte-specific deletion of *Pparα* promotes NASH in the context of obesity

**DOI:** 10.1101/488031

**Authors:** Marion Régnier, Arnaud Polizzi, Sarra Smati, Céline Lukowicz, Anne Fougerat, Yannick Lippi, Edwin Fouché, Frédéric Lasserre, Claire Naylies, Colette Bétoulières, Valentin Barquissau, Etienne Mouisel, Justine Bertrand-Michel, Aurélie Batut, Talal Al Saati, Cécile Canlet, Marie Tremblay-Franco, Sandrine Ellero-Simatos, Dominique Langin, Catherine Postic, Walter Wahli, Nicolas Loiseau, Hervé Guillou, Alexandra Montagner

## Abstract

**Objectives:** Peroxisome proliferator activated receptor α (PPARα) acts as a fatty acid sensor to orchestrate the transcription of genes coding for rate-limiting enzymes required for lipid oxidation in hepatocytes. Mice only lacking *Pparα* in hepatocytes spontaneously develop steatosis without obesity in aging. Altough steatosis is a benign condition it can develop into non alcoholic steatohepatitis (NASH), which may progress to irreversible damage, such as fibrosis and hepatocarcinoma. While NASH appears as a major public health concern worldwide, it remains an unmet medical need. Several drugs are being tested in clinical trials, including pharmacological agonists for the different PPAR isotypes. In current study, we investigated the role of hepatocyte PPARα in a preclinical model of steatosis.

**Methods/Results:** We have investigated the role of hepatocyte PPARα in a preclinical model of steatosis using High Fat Diet (HFD) feeding as a model of obesity in C57BL/6J male Wild-Type mice (*WT*), in whole-body (*Pparα^-/-^*) mice and in mice lacking *Pparα* in hepatocyte (*Pparα^hep-/-^*). We provide evidence that *Pparα* deletion in hepatocytes promotes NASH in mice fed an HFD. This enhanced NASH susceptibility occurs without development of glucose intolerance. Moreover, our data reveal that non-hepatocytic PPARα activity predominantly contributes to the metabolic response to HFD.

**Conclusion:** Taken together, our data support hepatocyte PPARα as being essential to the prevention of steatosis progression to NASH and that extra-hepatocyte PPARα activity contributes to whole-body lipid homeostasis.

**Highlights:**

- *Pparα* deletion in hepatocytes promotes steatosis and inflammation in HFD-induced obesity
- Hepatocyte-specific deletion of *Pparα* dissociates NAFLD from glucose intolerance in HFD-induced obesity
- Extrahepatic PPARα activity contributes to the metabolic response to HFD-induced obesity

## 1. Introduction

Non alcoholic fatty liver disease (NAFLD) has become a major public health concern worldwide [1]. NAFLD ranges from benign steatosis to non alcoholic steatohepatitis (NASH), which may progress to irreversible damage, such as fibrosis or hepatocarcinoma. The hallmark of NAFLD is an elevated level of neutral lipids, which accumulate as lipid droplets in hepatocytes [2]. Although the aetiology of the disease is not fully understood, it is strongly associated with obesity and type 2 diabetes (T2D). In human NAFLD, the fatty acids that accumulate in hepatocytes originate mostly from adipose tissue lipolysis and hepatic *de novo* lipogenesis [3]. In T2D, adipose tissue insulin resistance promotes lipolysis, whereas hyperglycaemia combined with hyperinsulinemia sustains hepatic *de novo* lipogenesis [4].

Given the burden of the NAFLD epidemic, identifying molecular players that can be targeted is a rather important issue [5,6]. Moreover, finding drugs that may be used to treat NASH and its progression to irreversible liver disease is a so far unmet medical need to be solved [7,8]. Among drugs currently being tested in clinical trials are a number of molecules that activate the peroxisome proliferator activated receptors (PPARs) [7,9]. Three PPAR isotypes are known (α, β/, and γ), and they are members of the nuclear receptor family, which act as fatty acid sensors that orchestrate transcription in response to a variety of endogenous ligands [9], such as fatty acids [10], fatty acid derivatives [11] and phospholipids [12]. Once activated by the binding of these lipids, PPARs may either induce or repress the expression of their specific target genes. PPARs are influential regulators of genes involved in metabolism in different tissues [13]. Therefore, several pharmacological agonists have been developed, tested on preclinical models of NAFLD [14,15], and are currently being either used or tested in clinical trials for the treatment of metabolic diseases [9], and especially NAFLD [15].

PPARα is the most abundant PPAR isotype in the healthy liver [16] and in hepatocytes, PPARα regulates the expression of thousands of genes and contributes to the remarkable metabolic flexibility of the liver [17–20]. PPARα is particularly active during suckling [21–23] and fasting [18,19,24–27], two conditions in which fatty acids are a preferred source of energy for the organism. PPARα is also expressed in many other tissues, including skeletal muscle [28], adipose tissues [29–32], intestine [33], heart [34], and kidney [35]. Germline deletion of *Pparα* renders mice susceptible for many metabolic defects including obesity [36], steatosis [36,37], and NASH [38,39], but not diabetes [40,41]. We have showed recently that a hepatocyte-specific deletion of *Pparα* induces spontaneous steatosis in aging mice and blunts fasting-induced ketogenesis [18,19]. Moreover PPARα is required for the expression of fibroblast growth factor 21 (FGF21) [42,43], a liver-derived hormone with many endocrine [44] and hepatoprotective effects [45,46].

In the present study, we evaluated the importance of hepatocyte PPARα in steatosis associated with diet-induced obesity. We provide evidence that in mice fed with a high fat diet (HFD), *Pparα* deletion in hepatocytes is sufficient to promote NAFLD. In addition, analysis of the hepatic transcriptome, lipidome, and metabolome, demonstrated that extrahepatic PPARα activity significantly contributes to metabolic homeostasis in response to HFD consumption.

## 2. Material and Methods

### 2.1. Mice

*In vivo* studies were conducted under the EU guidelines for the use and care of laboratory animals and they were approved by an independent ethics committee.

*Pparα^hep-/-^* animals were created at INRA’s rodent facility (Toulouse, France) by mating the floxed-*Pparα* mouse strain with C57BL/6J albumin-Cre transgenic mice (a gift from Prof. Didier Trono, EPFL, Lausanne, Switzerland) to obtain albumin-Cre+/−*Pparαflox/flox* mice (i.e., *Pparα^hep-/-^* mice). The *Pparα* deletion was confirmed with PCR and HotStar Taq DNA Polymerase (5 U/μl, Qiagen) using the following primers: forward: 5′-AAAGCAGCCAGCTCTGTGTTGAGC-3′ and reverse, 5′-TAGGTACCGTGGACTCAGAGCTAG-3′. The amplification conditions were as follows: 95°C for 15 min, followed by 35 cycles of 94°C for min, 65°C for 1 min, and 72°C for 1 min, and a final cycle of 72°C for 10 min. This reaction produced 450-bp, 915-bp, and 1070-bp fragments, which represented the *Pparα* sequence with an exon 4 deletion, the wild-type allele, and the floxed allele, respectively. The albumin-Cre allele was detected by PCR using the following primer pairs: CreU, 5′-AGGTGTAGAGAAGGCACTTAG-3′ and CreD, 5′-CTAATCGCCATCTTCCAGCAGG-3′; G2lox7F, 5′-CCAATCCCTTGGTTCATGGTTGC-3′ and G2lox7R, 5′-CGTAAGGCCCAAGGAAGTCCTGC-3′).

*Pparα*-deficient C57BL/6J mice (*Pparα^-/-^*) were bred at INRA’s transgenic rodent facility. Age-matched C57BL/6J (provided by Charles River) were acclimated to the local animal facility conditions prior to the experiment. Mouse housing was controlled for temperature and light (12-h light/12-h dark). All mice were placed in a ventilated cabinet at the specific temperature of 30°C (thermoneutrality) throughout the experiment. All animals used in these experiments were male mice.

### 2.2. Diet

*WT*, *Pparα^-/-^* and *Pparα^hep-/-^* mice were fed a standard diet (Safe 04 U8220G10R) until 8 weeks old, when the mice were fed a high fat diet (D12492, Research Diet) containing 60% calories from fat (lard), 20% calories from carbohydrates (7% sucrose) and 20% calories from proteins; or a chow diet (D12450J, Research Diet) containing 10% calories from fat, 70% calories from carbohydrates (7% sucrose) and 20% calories from protein during 12 weeks (until 20 weeks old). Experimental groups were designed as follows: *WT* CTRL, 8 mice; WT HFD, 8 mice; *Pparα^hep-/-^* CTRL, 10 mice; *Pparα^hep-/-^* HFD, 9 mice; *Pparα^-/-^* CTRL, 10 mice; *Pparα^-/-^* HFD, 10 mice.

### 2.3. Oral glucose tolerance test

Mice were fasted for 6 h and received an oral (2g/kg body weight) glucose load. Blood glucose was measured at the tail vein using an AccuCheck Performa glucometer (Roche Diagnostics) at −15, 0, 15, 30, 45, 60, 90, and 120 minutes.

### 2.4. Blood and tissue samples

Prior to sacrifice, blood was collected in EDTA-coated tubes (BD Microtainer, K2E tubes) from the submandibular vein. All mice were killed in a fed state. Plasma was collected by centrifugation (1500xg, 10min, 4°C) and stored at −80°C. Following killing by cervical dislocation, the organs were removed, weighted, dissected and used for histological analysis or snap-frozen in liquid nitrogen and stored at −80°C.

### 2.5. Gene expression

Total cellular RNA was extracted with Tri reagent (Molecular Research Center). Total RNA samples (2μg) were reverse-transcribed with the High-capacity cDNA Reverse Transcription Kit (Applied Biosystems) for real-time quantitative polymerase chain reaction (qPCR) analyses. The primers for Sybr Green assays are presented in Supplementary Table 1. Amplifications were performed on a Stratagene Mx3005P (Agilent Technology). The qPCR data were normalized to the level of the TATA-box binding protein (TBP) messenger RNA (mRNA) and analysed by LinRegPCR. Transcriptome profiles were determined by the Agilent SurePrint G3 Mouse GE v2 8×60K (Design 074809) according to the manufacturer’s instructions. Gene Ontology (GO) analysis of the KEGG categories was realized using string database and consists of an enrichment of biologically and functionally interacting proteins with a p-value ≤0.01. Microarray data and all experimental details are available in the Gene Expression Omnibus (GEO) database (accession GSE123354).

### 2.6. Histology

Formalin-fixed, paraffin-embedded liver tissue was sliced into 3-μm sections and stained with haematoxylin and eosin for histopathological analysis. The staining was visualized with a Leica microscope DM4000 B equipped with a Leica DFC450 C camera. The histological features were grouped into two categories: steatosis and inflammation. The steatosis score was evaluated according to Contos et al. [47]. Liver slices were assigned a steatosis score depending on the percentage of liver cells containing fat: Grade 0, no hepatocytes involved in any section; grade 1, 1% to 25% of hepatocytes involved; grade 2, 26% to 50% of hepatocytes involved; grade 3, 51% to 75% of hepatocytes involved and grade 4, 76% to 100% of hepatocytes involved. For the inflammation score, inflammatory foci were counted into 10 distinct areas at 200x for each liver slice. Values represent the mean of 10 fields/liver slice.

### 2.7. Biochemical analysis

Aspartate transaminase (AST), alanine transaminase (ALT), total cholesterol, LDL and HDL cholesterols were determined from plasma samples using a COBASMIRA+ biochemical analyser (Anexplo facility).

### 2.8. Analysis of liver neutral lipids

Tissue samples were homogenized in methanol/5 mM EGTA (2:1, v/v), and lipids (corresponding to an equivalent of 2 mg tissue) extracted according to the Bligh–Dyer method [48], with chloroform/methanol/water (2.5:2.5:2 v/v/v), in the presence of the following internal standards: glyceryl trinonadecanoate, stigmasterol, and cholesteryl heptadecanoate (Sigma). Triglycerides, free cholesterol, and cholesterol esters were analysed by gas-liquid chromatography on a Focus Thermo Electron system equipped with a Zebron-1 Phenomenex fused-silica capillary column (5 m, 0.25mm i.d., 0.25 mm film thickness). The oven temperature was programmed to increase from 200 to 350°C at 5°C/min, and the carrier gas was hydrogen (0.5 bar). The injector and detector temperatures were 315°C and 345°C, respectively.

### 2.9. Liver fatty acid analysis

To measure all hepatic fatty acid methyl ester (FAME) molecular species, lipids that corresponded to an equivalent of 1 mg of liver were extracted in the presence of the internal standard, glyceryl triheptadecanoate (2 μg) [49]. The lipid extract was transmethylated with 1 ml BF3 in methanol (14% solution; Sigma) and 1 ml heptane for 60 min at 80°C, and evaporated to dryness. The FAMEs were extracted with heptane/water (2:1). The organic phase was evaporated to dryness and dissolved in 50 μl ethyl acetate. A sample (1 μl) of total FAME was analysed by gas-liquid chromatography (Clarus 600 Perkin Elmer system, with Famewax RESTEK fused silica capillary columns, 30-m×0.32-mm i.d., 0.25-μm film thickness). The oven temperature was programmed to increase from 110°C to 220°C at a rate of 2°C/min, and the carrier gas was hydrogen (7.25 psi). The injector and detector temperatures were 225°C and 245°C, respectively.

### 2.10. Liver phospholipid and sphingolipid analysis

#### 2.10.1 Chemicals and reagents

The liquid chromatography solvent, acetonitrile, was HPLC-grade and purchased from Acros Organics. Ammonium formate (>99%) was supplied by Sigma Aldrich. Synthetic lipid standards (Cer d18:1/18:0, Cer d18:1/15:0, PE 12:0/12:0, PE 16:0/16:0, PC 13:0/13:0, PC 16:0/16:0, SM d18:1/18:0, SM d18:1/12:0) were purchased from Avanti Polar Lipids.

#### 2.10.2 Lipid extraction

Lipids were extracted from the liver (1 mg) as described by Bligh and Dyer in dichloromethane / methanol (2% acetic acid) / water (2.5:2.5:2 v/v/v). Internal standards were added (Cer d18:1/15:0, 16 ng; PE 12:0/12:0, 180 ng; PC 13:0/13:0, 16 ng; SM d18:1/12:0, 16 ng; PI 16:0/17:0, 30 ng; PS 12:0/12:0, 156.25 ng). The solution was centrifuged at 1500 rpm for 3 min. The organic phase was collected and dried under azote, then dissolved in 50 μl MeOH. Sample solutions were analysed using an Agilent 1290 UPLC system coupled to a G6460 triple quadripole spectrometer (Agilent Technologies). MassHunter software was used for data acquisition and analysis. A Kinetex HILIC column (Phenomenex, 50×4.6 mm, 2.6 μm) was used for LC separations. The column temperature was maintained at 40°C. Mobile phase A was acetonitrile and B was 10 mM ammonium formate in water at pH 3.2. The gradient was as follows: from 10% to 30% B in 10 min, 100% B from 10 to 12 min, and then back to 10% B at 13 min for 1 min to re-equilibrate prior to the next injection. The flow rate of the mobile phase was 0.3 ml/min, and the injection volume was 5μl. An electrospray source was employed in positive (for Cer, PE, PC, and SM analysis) or negative ion mode (for PI and PS analysis). The collision gas was nitrogen. Needle voltage was set at +4000 V. Several scan modes were used. First, to obtain the naturally different masses of different species, we analysed cell lipid extracts with a precursor ion scan at 184 m/z, 241 m/z, and 264 m/z for PC/SM, PI, and Cer, respectively. We performed a neutral loss scan at 141 and 87 m/z for PE and PS, respectively. The collision energy optimums for Cer, PE, PC, SM, PI, and PS were 25 eV, 20 eV, 30 eV, 25 eV, 45 eV, and 22 eV, respectively. The corresponding SRM transitions were used to quantify different phospholipid species for each class. Two MRM acquisitions were necessary, due to important differences between phospholipid classes. Data were treated with QqQ Quantitative (vB.05.00) and Qualitative analysis software (vB.04.00).

### 2.11. Metabolomic analyses by 1H nuclear magnetic resonance (NMR) spectroscopy

^1^H NMR spectroscopy was performed on aqueous liver extracts prepared from liver samples (50–75 mg) homogenized in chloroform/methanol/NaCl 0.9% (2/1/0.6) containing 0.1% butyl hydroxytoluene and centrifuged at 5000xg for 10 min. The supernatant was collected, lyophilized, and reconstituted in 600 μl of D2O containing 0.25 mM TSP [3-(trimethylsilyl)propionic-(2,2,3,3-d4) acid sodium salt] as a chemical shift reference at 0 ppm. All 1H NMR spectra were obtained on a Bruker DRX-600 Avance NMR spectrometer operating at 600.13 MHz for 1H resonance frequency using an inverse detection 5 mm 1H-13C-15N cryoprobe attached to a CryoPlatform (the preamplifier cooling unit). The 1H NMR spectra were acquired at 300 K with a 1D NOESY-presat sequence (relaxation delay – 90°-t-90°-tm-90°-acquisition). A total of 128 transients were acquired into a spectrum with 20 ppm width, 32 k data points, a relaxation delay of 2.0 s, and a mixing delay of 100 ms. All 1H spectra were zero-filled to 64 k points and subjected to 0.3 Hz exponential line broadening before Fourier transformation. The spectra were phase and baseline corrected and referenced to TSP (1H, d 0.0 ppm) using Bruker Topspin 2.1 software (Bruker GmbH, Karlsruhe, Germany). Multivariate analysis of metabolomic data was performed.

### 2.12. Statistical analysis

Data were analysed using R (http://www.r-project.org). Differential effects were assessed on log2 transformed data by performing analyses of variance, followed by an ANOVA test with a pooled variance estimate. p-values from ANOVA tests were adjusted with the Benjamini-Hochberg correction. p<0.05 was considered significant. Hierarchical clustering of gene expression data and lipid quantification data was performed with R packages, Geneplotter and Marray (https://www.bioconductor.org/). Ward’s algorithm, modified by Murtagh and Legendre, was used as the clustering method. All the data represented on the heat maps had adjusted p-values <0.05 for one or more comparisons performed with an analysis of variance.

## 3. Results

### 3.1. Hepatic and total *Pparα* deficiencies dissociate HFD-induced obesity and fatty liver from glucose intolerance

Male mice from different genotypes, namely wild-type (*WT*), germline *Pparα*-null (*Pparα^-/-^*) and hepatocyte-specific *Pparα*-null (*Pparα^hep-/-^*), were fed a low-fat diet (10% fat, CTRL) or a HFD (60% fat) at 8 weeks of age for 12 weeks at thermoneutrality (30°C). At the beginning of the experiment, the *Pparα^-/-^* mice were already significantly heavier than the *WT* and *Pparα^hep-/-^* mice (Figure 1A,B). All mice, independently of the genotype became overweight and gained approximately 15g in response to HFD consumption (Figure 1A). Moreover, unlike *WT* and *Pparα^hep-/-^* mice, *Pparα^-/-^* mice on CTRL diet also gained significant body weight. Therefore, *Pparα^-/-^* mice became more overweight than *Pparα^hep-/-^* and *WT* mice at thermoneutrality. In CTRL mice, oral glucose tolerance (OGTT) tested after 10 weeks of HFD feeding was similar regardless of the genotype (Figure 1C and D). In the HFD-fed groups, *WT* mice became glucose intolerant whereas *Pparα^hep-/-^* and *Pparα^-/-^* mice were protected against this intolerance (Figure 1C and D). These results are consistent with fasted glucose levels that increased in response to HFD only in *WT* mice, but not in *Pparα^hep-/-^* or *Pparα^-/-^* mice (Figure 1E). Therefore, HFD feeding leads to fasting hyperglycaemia and glucose intolerance in *WT* mice, but not in *Pparα^hep-/-^* and *Pparα^-/-^* mice.

**Figure 1.**
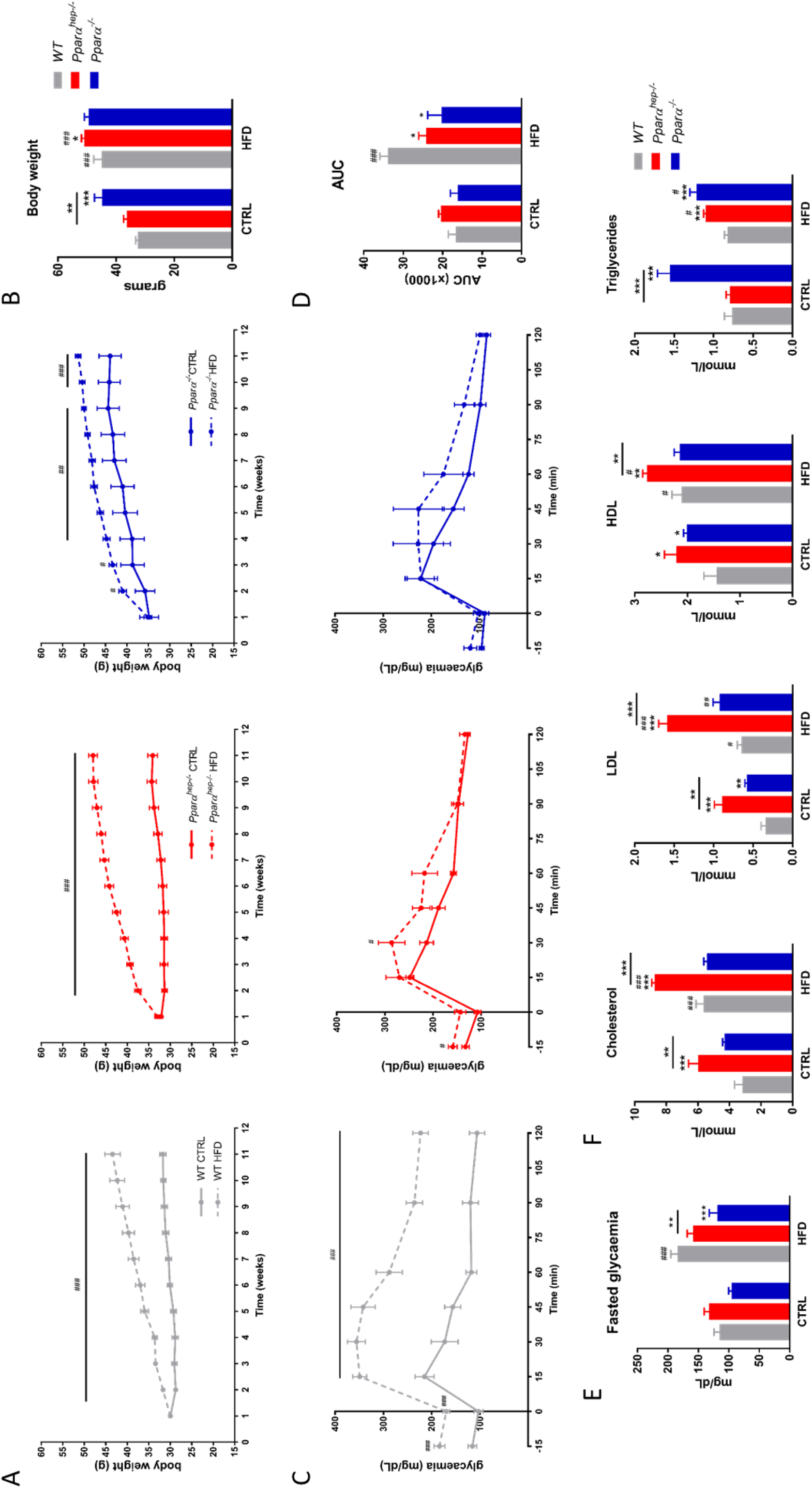
Hepatic and total *Pparα* deficiency does not promote glucose intolerance in HFD-induced obesity. *WT*, *Pparα^hep-/-^*, and *Pparα^-/-^* mice were fed a control diet (CTRL) or a HFD for 12 weeks at 30°C (thermoneutrality). (A) Body weight gain determined every week during the experiment. (B) Body weight at the end of the experiment. (C) Blood glucose measured during the oral glucose tolerance test (2g/kg of body weight). (D) Area under the curve obtained after the oral glucose tolerance test. (E) Quantification of fasted glycaemia. (F) Plasma cholesterol (total, HDL, and LDL) and triglyceride levels. Data represent mean ± SEM. #, significant diet effect and *, significant genotype effect. ^#^ or * p≤0.05; ## or ** p≤0.01; ^###^ or *** p≤0.001.

Different biochemical analyses were performed in plasma from fed animals (Figure 1F). Cholesterol, LDL-cholesterol, and HDL-cholesterol tended to increase in response to HFD diet in all three genotypes. However, we found that levels of the 3 lipid parameters were higher in the plasma of *Pparα^hep-/-^* mice than plasma from *Pparα^-/-^* mice. Triglycerides were elevated in *Pparα^-/-^* mice fed the CTRL diet and the HFD diet. Triglycerides were elevated in response to HFD only in *Pparα^hep-/-^* mice, but were lower in HFD-fed *Pparα^-/-^* mice compared to control diet-fed *Pparα^-/-^* mice. Taken together, our results show that both hepatocyte-specific and whole-body deletions of *Pparα* promote obesity dissociated from glucose intolerance in mice housed at thermoneutrality and fed a HFD.

### 3.2. Hepatic and total *Pparα* deficiencies promote liver steatosis, inflammation, and injury in HFD-induced obesity

Next, we performed histological analysis in order to investigate whether the lack of *Pparα* either globally or liver specific was associated with changes in liver integrity (Figure 2A). First, we observed that *Pparα^hep-/-^* and *Pparα^-/-^* mice developed steatosis upon CTRL diet feeding. In HFD, steatosis in *Pparα^hep-/-^* and *Pparα^-/-^* mice was much more severe than for WT mice, which is in agreement with their respective liver weight (Figure 2B). To better characterize liver injury, steatosis and inflammation scoring were performed (Figure 2C and E). The steatosis score revealed that *Pparα^hep-/-^* and *Pparα^-/-^* mice fed a HFD exhibited increasing lipid droplet deposition in the liver compared to *WT* mice (Figure 2C), which is confirmed by measurement of triglyceride liver content (Figure 2D). Inflammation scoring revealed that HFD did not significantly increase hepatic inflammation in WT mice (Figure 2E), contrarily to both *Pparα^-/-^* and *Pparα^hep-/-^* mice. In agreement with increased inflammation, plasma markers of liver injury (ALT and AST) were significantly increased in HFD fed *Pparα^-/-^* and *Pparα^hep-/-^* mice (Figure 2F).

**Figure 2.**
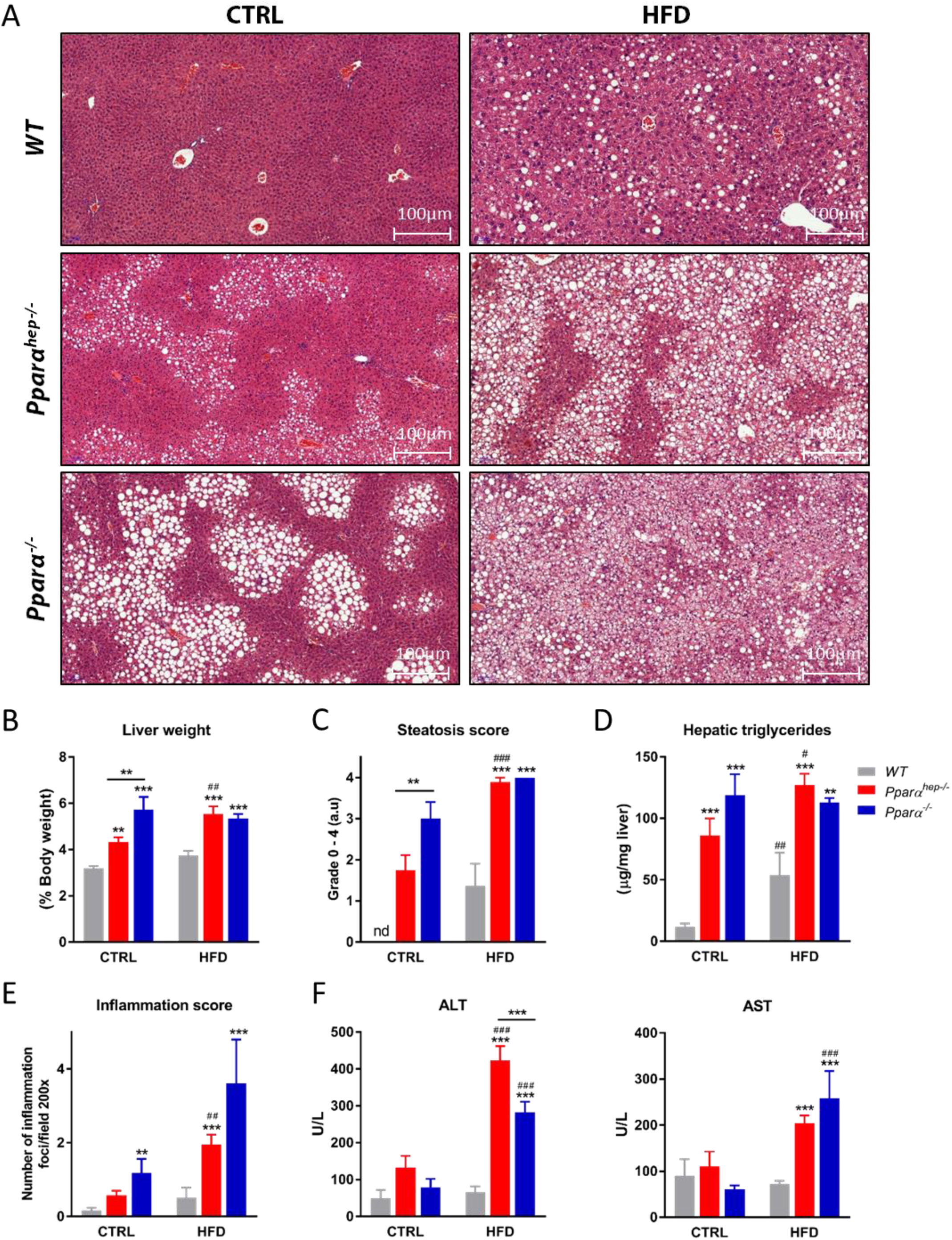
Hepatic and total *Pparα* deficiencies promote liver steatosis and inflammation in HFD induced obesity. *WT*, *Pparα^hep-/-^*, and *Pparα^-/-^* mice were fed a control diet (CTRL) or a HFD for 12 weeks at 30°C (thermoneutrality). (A) Representative pictures of Haematoxylin and Eosin staining of liver sections. Scale bars, 100μm. (B) Liver weight as a percentage of body weight. (C) Histological scoring of steatosis. (D) Quantification of hepatic triglycerides (E) Histological scoring of inflammation foci in 10 distinct areas at 200X. (F) Plasma ALT and AST. Data represent mean ± SEM. #, significant diet effect and *, significant genotype effect. ^#^ or * p≤0.05; ## or ** p≤0.01; ^###^ or *** p≤0.001.

### 3.3. Gene expression profile in *WT*, *Pparα^hep-/-^*, and *Pparα^-/-^* mice in response to HFD-induced obesity

Next, we evaluated the hepatic transcriptome expression pattern in response to HFD using microarrays. Overall, we identified a total of 8115 differentially expressed probes (corresponding to 7173 genes) sensitive to HFD feeding in at least one of the three genotypes (based on adjusted p-value; FDR<5%). Hierarchical clustering of probes highlighted 12 clusters showing specific gene expression profiles according to the experimental conditions (Figure 3A). Four of them (clusters 1, 2, 3 and 4) showed a typical transcriptomic pattern in *Pparα^-/-^* mice compared to *Pparα^hep-/-^* and *WT* mice regardless of diet. Interestingly, the liver transcriptome was also influenced by HFD in a genotype-specific manner (Figure 3A). The microarray analysis was validated by qPCR measurements of different genes (Figure 3B). The expression of *Vnn1*, encoding a liver-enriched oxidative stress sensor involved in the regulation of multiple metabolic pathways, was identified as an HFD-responsive gene specific to the *WT* mice. The expression of *Fmo3*, which produced trimethylamine N-oxide (TMAO) was identified as an HFD-induced gene specifically in *Pparα^-/-^* mice. The expression of collagen *Col1a1* is largely increased by HFD in the absence of hepatocyte-specific or whole-body *Pparα* and not in *WT* mice. Lastly, *Ppar-γ2* was identified as an HFD-responsive gene common to the three mouse genotypes. Significant overlaps occur between differentially expressed genes (DEGs) in the three genotypes (Figure 3C). Interestingly, only 387 DEGs were sensitive to HFD in all 3 genotypes at p<0.05, 232 at p<0.03 and 95 at p<0.01, mostly involved in metabolic regulations (Figure 3C and Supplementary Figure 1). This represents between 3 and 5% of all HFD-sensitive genes for which HFD regulation is strictly independent from hepatocyte or whole-body PPARα, therefore illustrating the critical role of the nuclear receptor in the liver response to a hyperlipidic diet.

**Figure 3.**
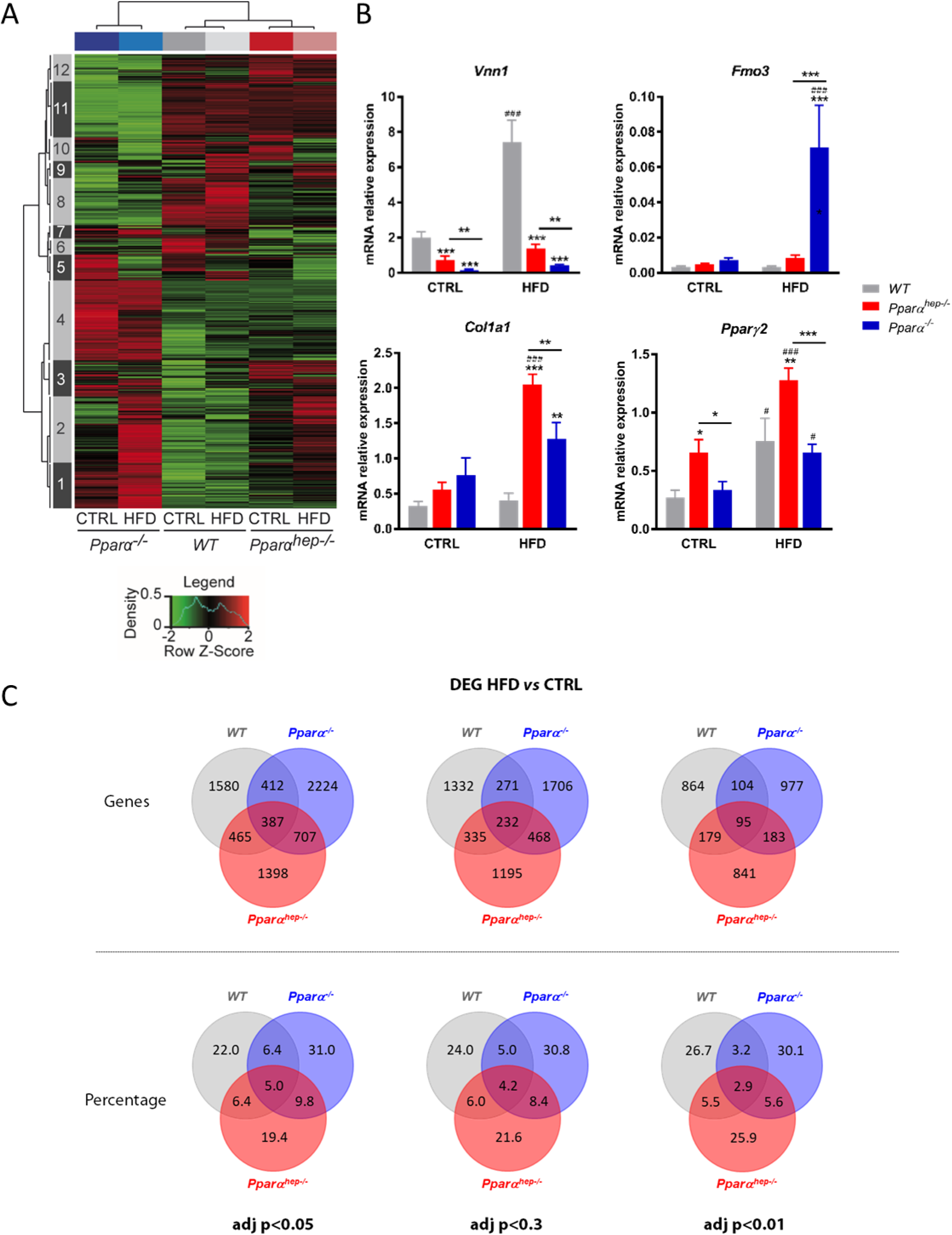
Analysis of the liver transcriptome in *WT*, *Pparα^hep-/-^*, and *Pparα^-/-^* mice in response to HFD. *WT*, *Pparα^hep-/-^*, and *Pparα^-/-^* mice were fed a control diet (CTRL) or a HFD for 12 weeks at 30°C (thermoneutrality). A transcriptomic analysis performed with liver samples from *WT*, *Pparα^hep-/-^*, and *Pparα^-/-^* exposed or not exposed to HFD (n=8 mice/group) revealed 8860 differentially regulated probes (FDR<5%). (A) Heat map of microarray expression data from 7173 regulated genes. Red and green indicate values above and below the mean averaged centred and scaled expression values (Z-score), respectively. Black indicates values close to the mean. According to the probe clustering (left panel), 12 gene clusters exhibited specific gene expression profiles. (B) Relative hepatic expression of *Vnn1*, *Fmo3, Col1a1* and *Pparγ2* quantified by qPCR. Data represent mean ± SEM. #, significant diet effect and *, significant genotype effect. ^#^ or * p≤0.05; ## or ** p≤0.01; ^###^ or *** p≤0.001. (C) Venn diagrams comparing the number (top) and the percentage (down) of genes significantly regulated under HFD in the livers of *WT*, *Pparα^hep-/-^*, and *Pparα^-/-^* mice at adjusted p-value<0.05, <0.03 and <0.01, respectively.

### 3.4. PPARα-dependent changes in hepatic gene expression profiles in response to HFD induced obesity

A large group of DEGs include 922 genes significantly up-regulated and 857 significantly down-regulated by HFD feeding only in *WT* mice (Figure 4A). Examples of these genes include well-established PPARα targets, such as *Cyp4a14*, *Acot3*, *Acot2*, and *Fitm1* (Figure 4B), and eight categories of genes involved in metabolism (Figure 4C) with up-regulated expression in response to a HFD only in *WT* mice. However, we also identified four KEGG categories down-regulated in response to HFD specifically in *WT* mice. These categories relate to RNA transport, ribosome activity and protein processing (Figure 4D). The venn diagram (Figure 3C) also reveals a large group of DEGs modulated only in *Pparα^hep-/-^* mice fed a HFD compared to CTRL diet which encompassed 630 and 865 significantly up or down-regulated genes, respectively (Supplementary Figure 2). Figure 3C also defines a third large group of DEGs including 1143 and 1257 genes significantly up-regulated and down-regulated, respectively, by HFD feeding only in *Pparα^-/-^* mice (Supplementary Figure 3). This suggests that the hepato-specific and whole-body deletions of *Pparα* have distinct and specific consequences in the hepatic response to HFD-induced obesity.

### 3.5. Hepatocyte PPARα prevents liver inflammatory gene expression in response to HFD

Next, we questioned whether deletion of *Pparα* in hepatocytes only or in the whole-body induces some similar responses in HFD-induced obesity. We identified a group of DEGs including 337 and 349 genes significantly up-regulated or down-regulated, respectively, by HFD feeding in both *Pparα^-/-^* and *Pparα^hep-/-^* mice (Figure 5A). Gene category analysis did not reveal any functions related to the 349 down-regulated genes by HFD feeding in *Pparα^-/-^* or *Pparα^hep-/-^* mice. However, gene category analysis highlighted the functions related to the 337 genes significantly up-regulated by HFD feeding in both *Pparα^-/-^* and *Pparα^hep-/-^* mice (Figure 5B), suggesting that these genes are negatively regulated by PPARα Most of these categories relate to the inflammatory process, including the NF-kappa B, TNF, and TLR signalling pathways. We selected the genes directly related to these pathways using the KEGG database and the gene database network (Supplementary Figure 4) and confirmed a marked up-regulation of genes belonging to NF-kappa B, TNF, and TLR in the hepatocyte-specific or whole-body absence of *Pparα* (Figure 5C), in accordance with inflammatory markers measured (Figure 2E).

**Figure 4.**
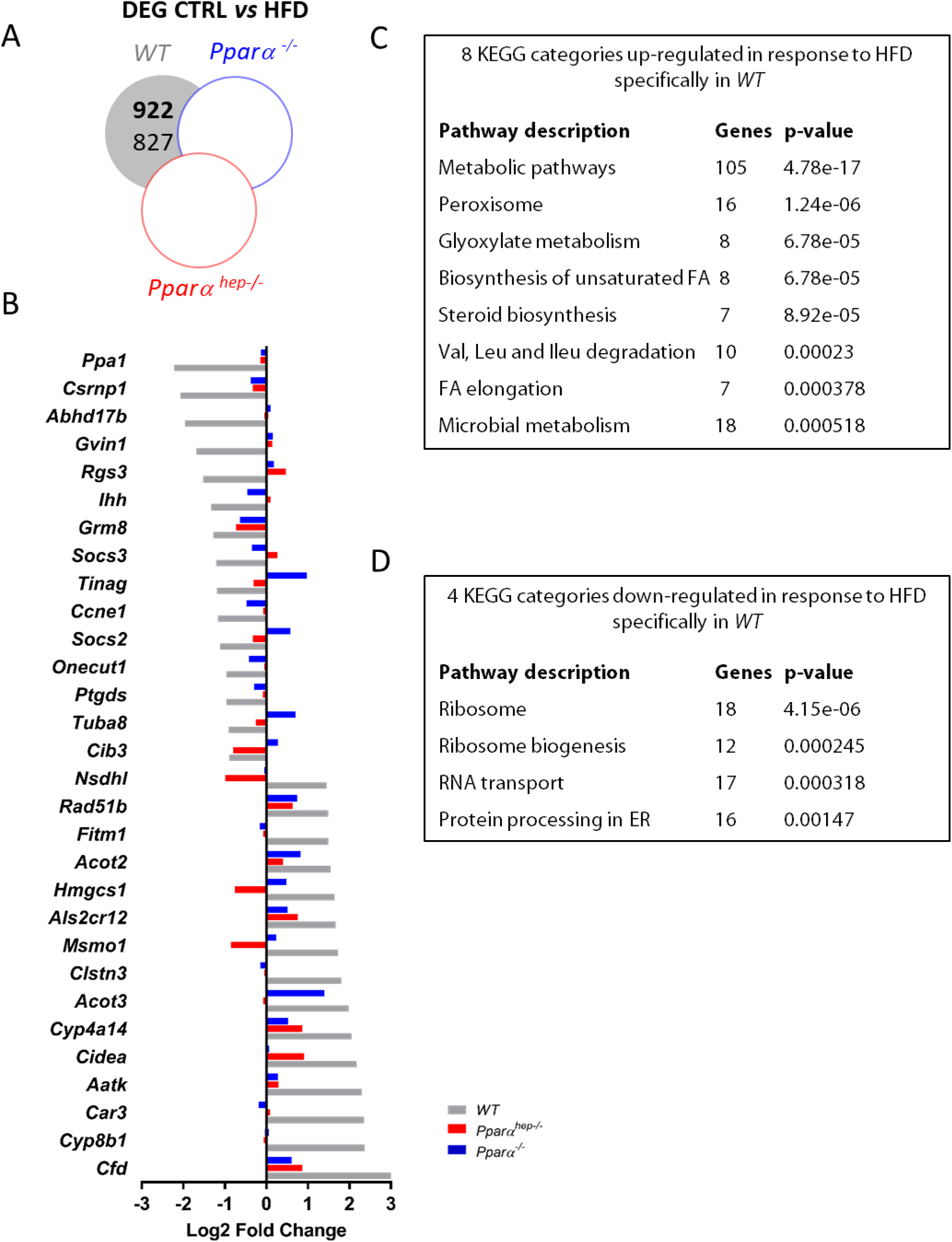
PPARα-dependent changes in hepatic gene expression profiles in response to HFD. *WT*, *Pparα^hep-/-^*, and *Pparα^-/-^* mice were fed a control diet (CTRL) or a HFD for 12 weeks at 30°C (thermoneutrality). (A) Venn diagram presenting the number of hepatic genes over-expressed (bold) and down-regulated (regular) in response to HFD in *WT*, *Pparα^hep-/-^*, *Pparα^-/-^* mice (FDR<5%) (B) Grey bars represent the top 15 specifically induced and repressed genes between *WT* exposed to CTRL diet and *WT* exposed to HFD. Red and blue bars represent the profile in *Pparα^hep-/-^* and *Pparα^-/-^* mice, respectively. (C) Gene Ontology (GO) enrichment analysis (p≤0.01) of KEGG categories based on functional interactions specifically down-regulated in *WT* mice fed a HFD using the String database. (D) Gene Ontology (GO) enrichment analysis (adjusted p-value; p≤0.01) of KEGG categories (based on functionally interactions) up-regulated in *WT* mice fed a HFD the using string database.

**Figure 5.**
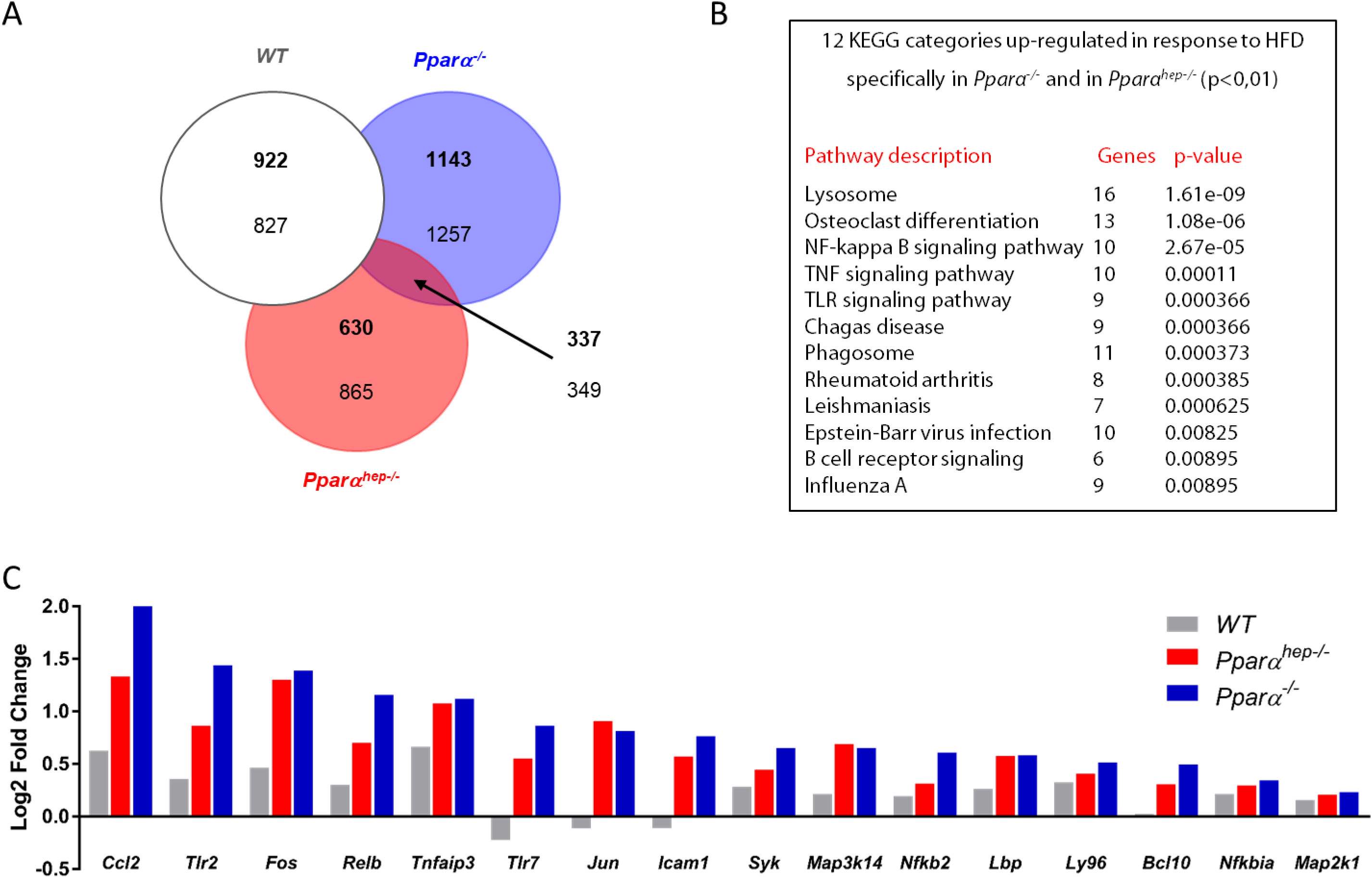
Hepatocyte PPARα prevents liver inflammatory gene expression in response to HFD. *WT*, *Pparα^hep-/-^*, and *Pparα^-/-^* mice were fed a control diet (CTRL) or a HFD for 12 weeks at 30°C (thermoneutrality). (A) Venn diagram highlighting the number of hepatic genes over-expressed (bold) and down-regulated (regular) in response to HFD specifically in both *Pparα^hep-/-^* and *Pparα^-/-^* mice (FDR<5%). (B) Gene Ontology (GO) enrichment analysis (adjusted p≤0.01) of KEGG categories specifically up-regulated in *Pparα^hep-/-^* and *Pparα^-/-^* mice fed a HFD. (C) Gene expression profile of genes identified as being ninvolved in NF-kappa B, TNF and TLR signalling pathways in KEGG.

### 3.6. Metabolic and lipidomic profiling of PPARα-dependent regulation of hepatic homeostasis in response to HFD

We performed unbiased hepatic metabolomic profiling of aqueous metabolites using proton nuclear magnetic resonance (^1^H-NMR). We used a projection to latent structures for discriminant analysis (PLS-DA) to investigate whether there was a separation between experimental groups of observations. A valid and robust PLS-DA model was obtained that discriminated HFD-fed *Pparα^-/-^* mice from all other groups (Supplementary Figure 5), further supporting the role of non-hepatocytic PPARα activity in hepatic homeostasis in response to HFD.

We also performed a targeted analysis of 75 lipid species including neutral lipids (cholesterol, cholesterol esters, and triglycerides), phospholipids, and sphingolipids. The relative abundance of each species in the livers of *WT*, *Pparα^-/-^*, and *Pparα^hep-/-^* mice fed with either of the two diets (CTRL and HFD) was evaluated to determine the contribution of hepatocyte and whole body PPARα activity to hepatic lipid homeostasis. The results are presented as a heatmap with hierarchical clustering (Figure 6A), in which we observed that the samples first clustered according to the diet, demonstrating that HFD-feeding was the main discriminating factor for hepatic lipid content. We identified four main clusters of lipids with distinct profiles relative to the different experimental conditions. Lipids in cluster 1, such as the ceramides d18:1/C18:1, d18:1/C18:0, and d18:1/C26:0 (Figure 6B), exhibit increased relative abundance in HFD-fed *Pparα^-/-^* mice, suggesting that extra-hepatocytic PPARα contributes predominantly to lipid remodelling during HFD-feeding. Cluster 1 also contains linoleic acid (C18:2n-6), which exhibits increased abundance in HFD-fed *Pparα^-/-^* mice, but also in HFD-fed *Pparα^hep-/-^* mice. Lipids in cluster 2, such as the phospholipids PC36:3, PC28:6, PE38:4, and triglyceride TG C57 (Figure 6C), are less abundant in HFD *Pparα^-/-^* mice. Lipids in cluster 3, such as the palmitoleic acid (C16:1n-7) and PE32:1 (Figure 6D), are less abundant in HFD mice from the three genotypes. Lipids in cluster 4 (Figure 6E), such as the polyunsaturated fatty acids C20:4n-6 and C22:5n-3 are more abundant in WT mice from the CTRL diet group and reduced in the livers of mice fed a HFD.

**Figure 6.**
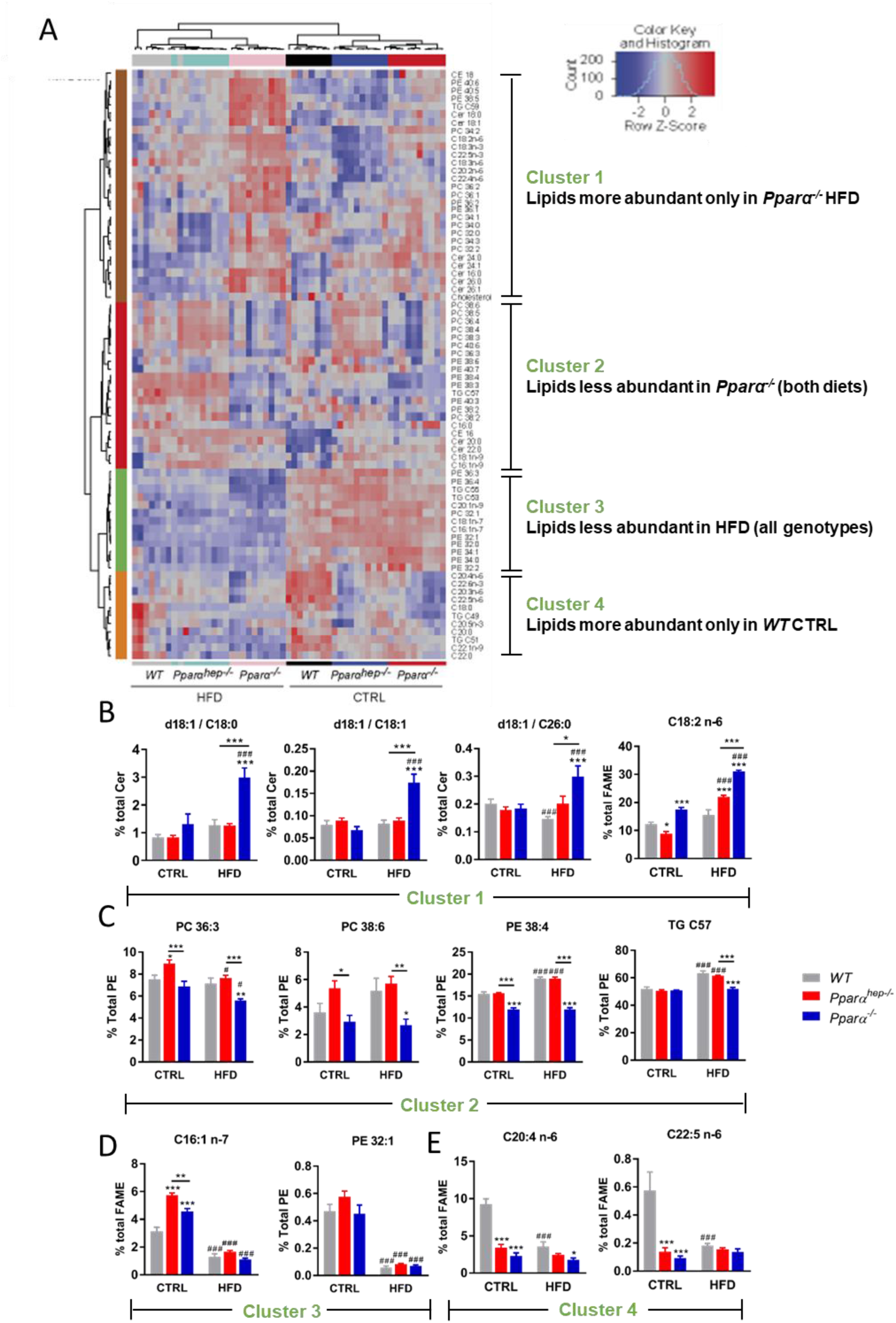
PPARα-dependent regulation of hepatic lipid homeostasis in response to HFD. *WT*, *Pparα^hep-/-^*, and *Pparα^-/-^* mice were fed a control diet (CTRL) or a HFD for 12 weeks at 30°C (thermoneutrality). (A) Heat map of data from hepatic lipid profiling in *WT*, *Pparα^hep-/-^*, and *Pparα^-/-^* mice exposed to HFD. Hierarchical clustering is also shown and defines four main lipid clusters. Representation of characteristic lipid species defining cluster 1 (B), 2 (C), 3 (D), and 4 (E). Data represent mean ± SEM. Data represent mean ± SEM. #, significant diet effect and *, significant genotype effect. ^#^ or * p≤0.05; ## or ** p≤0.01; ^###^ or *** p≤0.001. Cer: ceramide; SM: sphingomyelin; TG: triglyceride; PC: phosphatidyl choline; PE: phosphatidyl ethanolamine; CE: cholesterol ester.

Overall, this lipidomic profiling highlights that the hepatic lipidome depends both on the genotype and diet. Therefore, both hepatic *Pparα* and whole body *Pparα* deletions result in a specific lipidomic response to HFD feeding. However, whole body *Pparα* deficiency has a stronger influence on the effect of HFD-induced obesity on liver metabolic homeostasis.

## 4. Discussion

NAFLD is the hepatic manifestation of the obesity epidemic and represents a major public health issue worldwide [50]. NAFLD ranges from benign steatosis to NASH and may promote liver fibrosis and cancer. Therefore, there is a great interest in drugs that could be used to cure NAFLD or reverse NASH before it promotes irreversible damage [7]. PPARα, and other PPAR isotypes represent targets currently being tested in clinical trials [9,51,52]. PPARα is a ligand-activated nuclear receptor that plays a key role in the regulation of metabolic homeostasis by modulating the expression of rate-limiting enzymes involved in fatty acid degradation [17–19]. Preclinical studies in *Pparα* null mice have shown that PPARα protects from steatosis [18,53] and NASH. Moreover, several clinical lines of evidence indicate that PPARα is also influential in human NASH [54].

Most studies performed *in vivo* in *Pparα* null mice have suggested that the mechanisms by which PPARα protects from steatosis and NASH involve its ability to transactivate genes required for fatty acid catabolism and to repress a number of inflammatory genes [38,39]. Because PPARα is expressed in many cell types and tissues with high fatty acid oxidation activity [28–30,33], it is interesting to define *in vivo* the specific contribution of hepatocytic PPARα in preventing NAFLD.

When fed a regular diet, *Pparα^-/-^* mice are steatotic and overweight as previously reported [25,37]. When fed a HFD, they become even more steatotic and develop further liver inflammation and NASH injury [38,39]. Importantly, although *Pparα*-null mice develop steatosis, they do not exhibit reduced glucose tolerance compared to *WT* mice [41]. The data we obtained in *Pparα^hep-/-^* mice fed a HFD indicate that the deletion of *Pparα* in hepatocytes is sufficient to promote steatosis and marked signs of fibrosis and inflammation. As in *Pparα*-null mice, despite enhanced steatosis compared to HFD-fed *WT* mice, HFD-fed *Pparα^hep-/-^* mice do not exhibit enhanced glucose intolerance. Therefore, hepatocyte-specific *Pparα* deletion promotes steatosis and liver inflammation. Interestingly, it also dissociates steatosis from glucose intolerance as observed in *Pparα^-/-^* mice [40,41].

The mechanisms involved in the susceptibility to steatosis and protection from glucose intolerance likely involve the well-established role of PPARα in the control of fatty acid transport and degradation [17]. The mechanisms by which hepatocyte *Pparα* deficiency promotes NASH likely involve lipotoxic fat accumulation, including linoleic acid (C18:2n-6), which was recently identified as promoting NAFLD etiology *in vivo* [55]. Moreover, we extend previous observations and confirm the role of hepatocyte PPARα in repressing the expression of inflammatory genes, such as those involved in the NF-kappa B pathway [56,57].

This study not only shows a specific effect of hepatocyte PPARα activity, but also identifies several roles of non-hepatocytic PPARα. First, we confirmed our previous observations that, unlike *Pparα^hep-/-^* mice, *Pparα^-/-^* mice gain weight when fed a regular diet [18]. Moreover, because the current observations were made at thermoneutrality, this weight gain is not likely due to defective PPARα expression and activity in the brown adipose tissue. In addition, by combining different large or medium scale analyses of the liver transcriptome and metabolome, we showed that the response to HFD was different in *Pparα^-/-^* mice compared to *Pparα^hep-/-^* mice, showing that extra-hepatocyte PPARα mediates at least a part of the adaptive response to HFD. These data are in agreement with a recent study providing evidence that extrahepatic PPARα such as in skeletal muscle and heart, may contribute to whole-body fatty acid homeostasis [58].

Taken together, our data demonstrate that hepatocyte-specific deletion of *Pparα* promotes steatosis and NASH in HFD-induced obesity and provide further pre-clinical evidence that hepatocyte PPARα is a relevant direct target in NAFLD. Our data also suggest that extra-hepatic PPARα plays a major homeostatic role in the control of the metabolic response to HFD. Further research is required to investigate in which organ, other than liver, PPARα regulates lipid metabolism in health and disease.

## Acknowledgments

We thank all members of the EZOP staff for their careful help from the early start of this project. We thank Léa Morra-Charrot and Laurent Monbrun for their excellent work on plasma biochemistry. We thank the staff from the Genotoul: Anexplo, GeT-TRiX and Metatoul-Lipidomic facilities. The authors wish to thank Pr Daniel Metzger, Pr Pierre Chambon (IGBMC, Illkirch, France) and the staff of the Mouse Clinical Institute (Illkirch, France) for their critical support in this project. We thank Pr Didier Trono (EPFL, Lausanne, Switzerland) for providing the Albumin-Cre mice. We thank Pr Bart Staels for constructive discussions. We thank Dr Thierry Pineau for constructive discussions from the early start of this project and for providing us with PPAR-null mice.

## Figure legends

**Supplementary Table 1. Oligonucleotide sequences for real-time qPCR.** Oligonucleotides (Sigma-Aldrich) were designed in Primer Express 2.0 software (Applied Biosystems). Couples of primers have a Tm of 60°C, exhibit no amplification with genomic DNA, and the derivative from each dissociation curve of amplicons from cDNA exhibits a single specific peak.

**Supplementary Figure 1. Common changes in the hepatic gene expression profiles of *WT*, *Pparα^hep-/-^*, *Pparα^-/-^* mice in response to HFD.** (A) Venn diagram presenting the number of overlapping hepatic genes up-regulated (bold) and down-regulated (regular) in response to HFD in *WT*, *Pparα^hep-/-^*, and *Pparα^-/-^* mice (adjusted p≤0.05) (B) Grey bars represent the top 15 induced and repressed genes specifically in *WT* mice exposed to CTRL diet vs HFD. Red and blue bars represent the HFD-induced responses in *Pparα^hep-/-^* and *Pparα^-/-^*, respectively. (C) Analysis of KEGG categories up-regulated in *WT*, *Pparα^hep-/-^* and *Pparα^-/-^* mice fed a HFD vs. CTRL diet (p≤0.01). (D) Analysis of KEGG categories down-regulated in *WT*, *Pparα^hep-/-^* and *Pparα*^-/-^ mice fed a HFD (p≤0.01).

**Supplementary Figure 2. Specific changes in liver gene expression profiles in *Pparα^hep-/-^* mice fed a HFD.** (A) Venn diagram presenting the number of hepatic genes up-regulated (bold) and down-regulated (regular) in response to HFD in *WT*, *Pparα^hep-/-^* and *Pparα^-/-^* mice (adjusted p-value; p≤0.05) (B) Red bars represent the top 15 induced and repressed genes in *Pparα^hep-/-^* mice fed a HFD vs CTRL diet. Grey bars represent the gene expression profile in WT mice and blue bars the gene expression profile in *Pparα^-/-^* mice (C) Analysis of KEGG categories specifically down-regulated in *Pparα^hep-/-^* mice fed a HFD (p≤0.01).

**Supplementary Figure 3. Specific changes in liver gene expression profiles in *Pparα^-/-^* mice fed a HFD.** (A) Venn diagram presenting the number of hepatic genes up-regulated (bold) and down-regulated (regular) in response to HFD in *WT*, *Pparα^hep-/-^* and *Pparα^-/-^* mice (adjusted p≤0.05). (B) Blue bars represent the top 15 induced and repressed genes in *Pparα^-/-^* mice fed a HFD vs. CTRL diet. Grey bars represent the gene expression profile in *WT* mice and red the gene expression profile in *Pparα^hep-/-^* mice (C) Analysis of KEGG categories specifically up-regulated in *Pparα^-/-^* mice fed a HFD (p≤0.001).

**Supplementary Figure 4. Identification of genes related to NF-kappa B, TNF, and TLR signalling pathways.** The predicted gene-gene interaction network (Search Tool for the Retrieval of Interacting Genes/StringV10) among genes significantly induced in *Pparα^hep-/-^*, *Pparα^-/-^* mice (adjusted p≤0.05) fed a HFD. Genes coloured in red, blue, and green belong to the GO categories “NF-kappa B signalling pathway”, “TNF signaling pathway”, and “TLR signalling pathway”.

**Supplementary Figure 5. Liver metabolome discriminates whole-body *Pparα* deletion from hepatocyte-specific PPARα deletion in mice fed a HFD.** Two-dimensional PLS-DA score plot of liver extract integrated 1H-NMR spectra. Each dot represents an observation (animal), projected onto first (horizontal axis) and second (vertical axis) PLS-DA variables.

