## Supplemental Table 1 for "Hepatocyte-specific deletion of *Pparα* promotes NASH in the context of obesity"

| Gene | NCBI Refseq | Forward primer (5'-3') | Reverse primer (5'-3') |
|---|---|---|---|
| *Col1a1* | NM_007742 | GGCTCCTGCTCCTCTTAGGG | TCGGGTTTCCACGTCTCAC |
| *Fmo3* | NM_008030 | AAGAAAGGAAGACAAAGAAAAGGCA | AGCTCCAATGATGGCCACTT |
| *Ppar-γ2* | NM_011146 | GATGCACTGCCTATGAGCACTT | GAATGGCATCTCTGTGTCAACC |
| *Vnn1* | NM_011704 | ATGAGGTTTATGCCTTTGGAGC | CCACAGGTGCGTAAATTGGTAG |
