## Supplemental Figure 1 for "Hepatocyte-specific deletion of *Pparα* promotes NASH in the context of obesity"

A

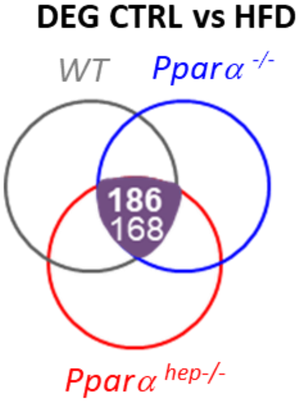


B

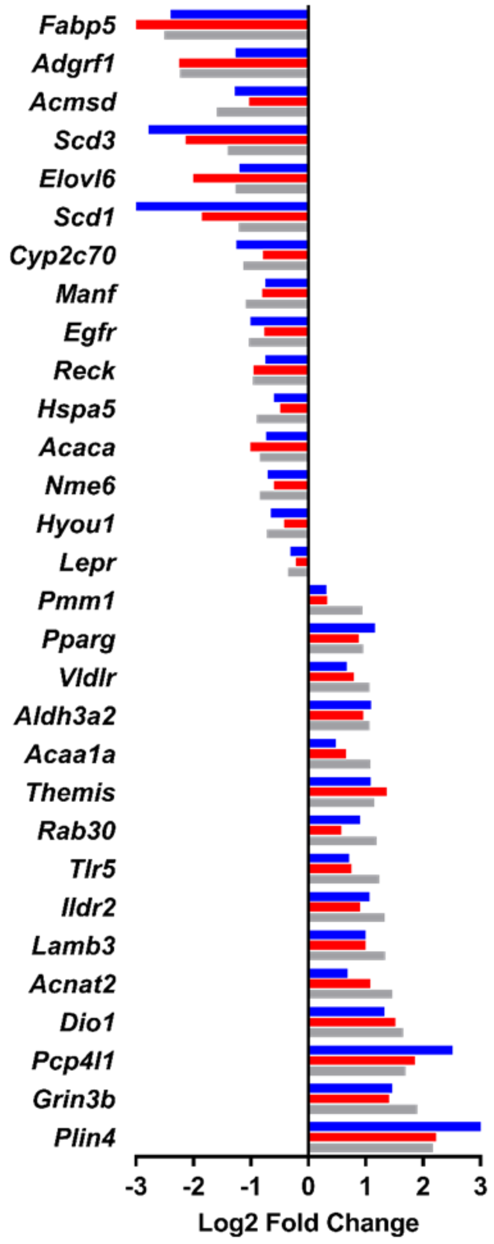


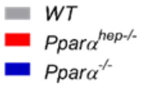


C

15 KEGG categories up-regulated in response to HFD in *WT*, *Pparα*$^{hep-/-}$ and *Pparα*$^{-/-}$

| Pathway description | Genes | p-value |
|---|---|---|
| Metabolic pathways | 37 | 7.35e-12 |
| Fatty acid degradation | 10 | 6.4e-11 |
| Butanoate metabolism | 7 | 3.48e-08 |
| Fatty acid metabolism | 8 | 1.35e-07 |
| Propanoate metabolism | 6 | 3.65e-06 |
| Microbial metabolism | 10 | 1.61e-05 |
| Ketone bodies metabolism | 4 | 1.81e-05 |
| Carbon metabolism | 7 | 0.000378 |
| PPAR signaling pathway | 6 | 0.000579 |
| Peroxisome | 6 | 0.000654 |
| beta-Alanine metabolism | 4 | 0.00161 |
| Pyruvate metabolism | 4 | 0.00497 |
| Tryptophan metabolism | 4 | 0.00554 |
| Renin-angiotensin system | 3 | 0.00604 |
| Lysine degradation | 4 | 0.00858 |

D

3 KEGG categories downregulated in response to HFD in *WT*, *Pparα*$^{hep-/-}$ and *Pparα*$^{-/-}$

| Pathway description | Genes | p-value |
|---|---|---|
| Protein processing in ER | 17 | 3.27e-13 |
| Protein export | 8 | 3.24e-10 |
| Fatty acid metabolism | 5 | 0.002 |
