## Supplemental Figure 2 for "Hepatocyte-specific deletion of *Pparα* promotes NASH in the context of obesity"

A

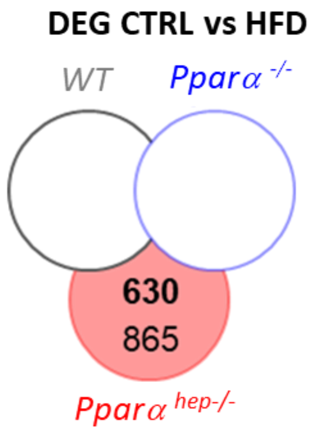


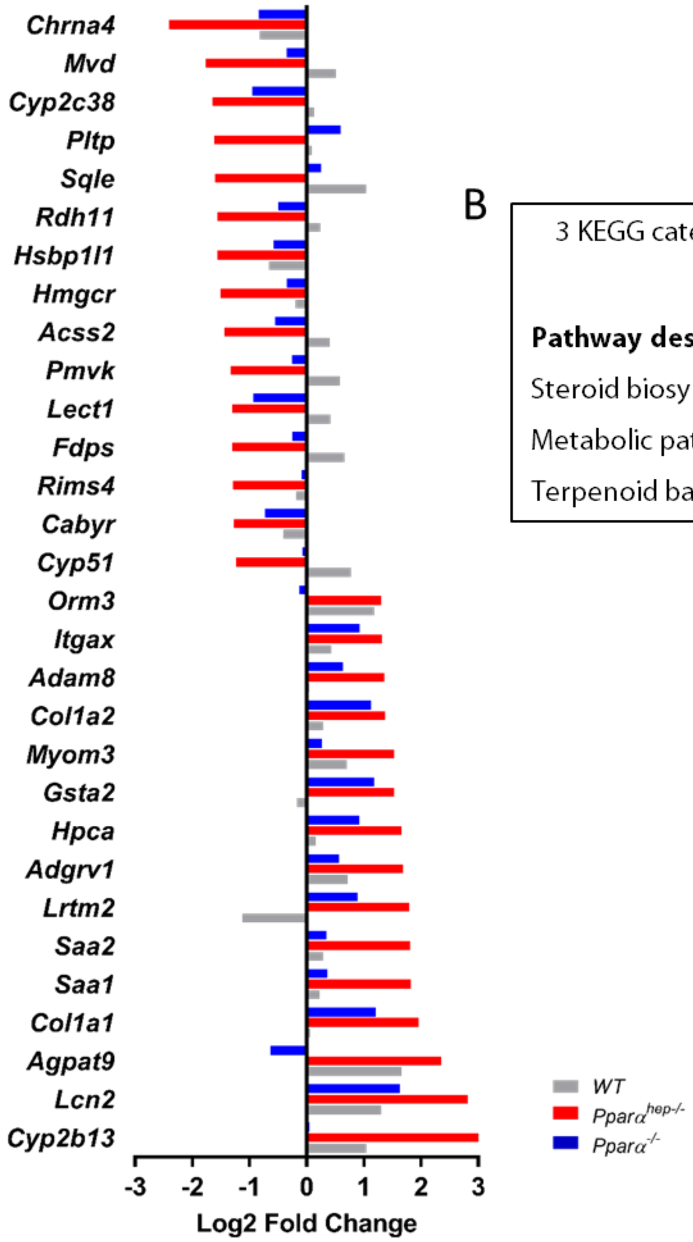


B

3 KEGG categories down-regulated in response to HFD specifically in *Pparα*$^{hep-/-}$

| Pathway description | Genes | p-value |
|---|---|---|
| Steroid biosynthesis | 10 | 2.91e-09 |
| Metabolic pathways | 63 | 2.43e-05 |
| Terpenoid backbone biosynthesis | 7 | 6.4e-05 |
