## Supplemental Figure 3 for "Hepatocyte-specific deletion of *Pparα* promotes NASH in the context of obesity"

A

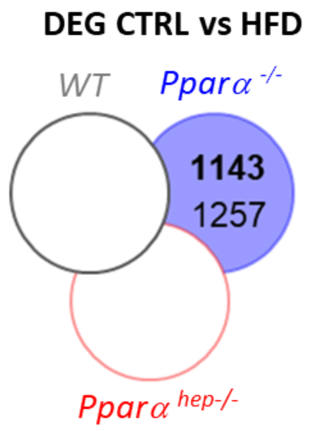


B

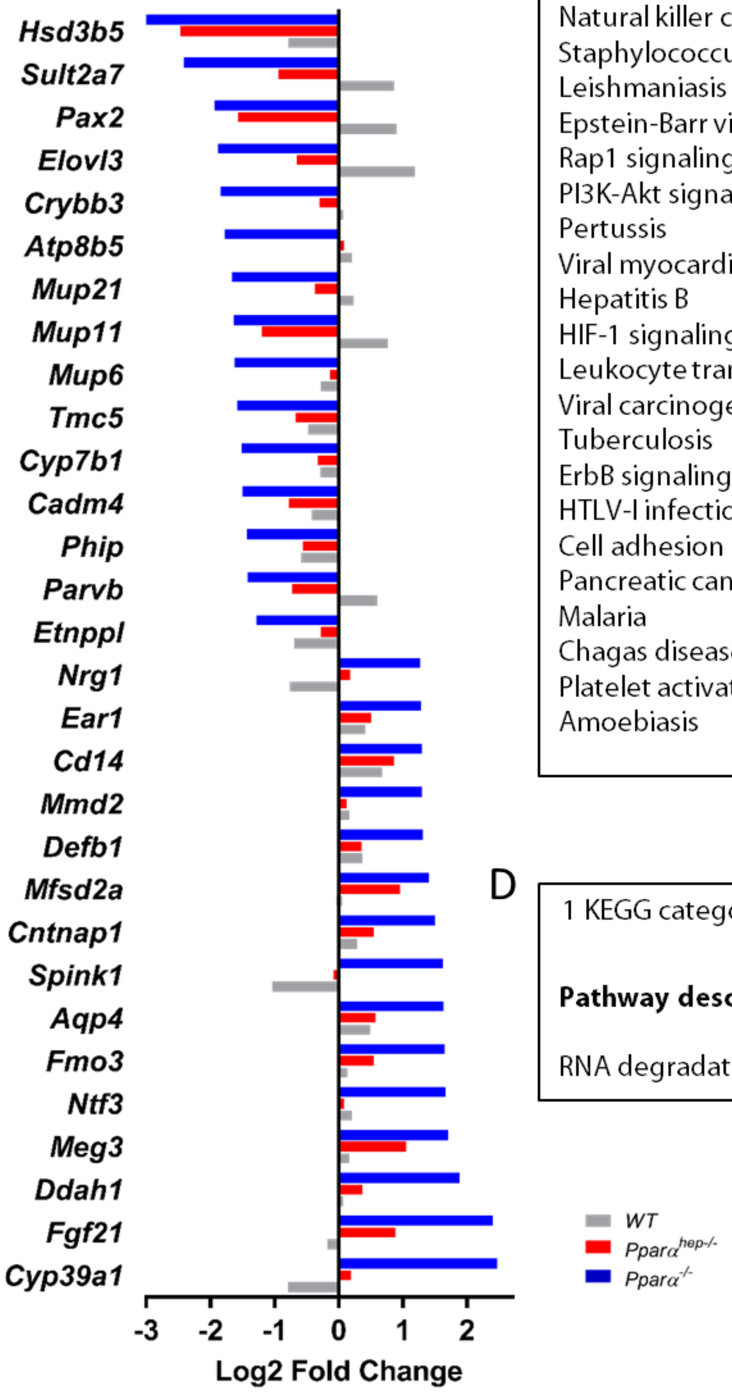


C

28 KEGG categories up-regulated in response to HFD specifically in *Pparα*$^{-/-}$

| Pathway description | Genes | p-value |
|---|---|---|
| Osteoclast differentiation | 26 | 5.06e-10 |
| Fc gamma R-mediated phagocytosis | 22 | 5.06e-10 |
| Ras signaling pathway | 31 | 1.63e-07 |
| Fc epsilon RI signaling pathway | 17 | 1.63e-07 |
| MAPK signaling pathway | 32 | 7.05e-07 |
| Chemokine signaling pathway | 25 | 3.32e-06 |
| Toll-like receptor signaling pathway | 18 | 4.79e-06 |
| Natural killer cell mediated cytotoxicity | 19 | 5.8e-06 |
| Staphylococcus aureus infection | 12 | 1.92e-05 |
| Leishmaniasis | 13 | 4.59e-05 |
| Epstein-Barr virus infection | 24 | 5.58e-05 |
| Rap1 signaling pathway | 24 | 0.000106 |
| PI3K-Akt signaling pathway | 33 | 0.000106 |
| Pertussis | 13 | 0.000106 |
| Viral myocarditis | 13 | 0.000106 |
| Hepatitis B | 19 | 0.000114 |
| HIF-1 signaling pathway | 16 | 0.000124 |
| Leukocyte transendothelial migration | 17 | 0.000144 |
| Viral carcinogenesis | 22 | 0.000314 |
| Tuberculosis | 20 | 0.000329 |
| ErbB signaling pathway | 13 | 0.000488 |
| HTLV-I infection | 26 | 0.000497 |
| Cell adhesion molecules (CAMs) | 18 | 0.000633 |
| Pancreatic cancer | 11 | 0.000633 |
| Malaria | 9 | 0.000699 |
| Chagas disease | 14 | 0.000785 |
| Platelet activation | 16 | 0.000827 |
| Amoebiasis | 15 | 0.000998 |

D

1 KEGG category down-regulated in response to HFD specifically in *Pparα*$^{-/-}$

| Pathway description | Genes | p-value |
|---|---|---|
| RNA degradation | 12 | 0.00941 |
