## Supplementary figures and images for "Hepatocyte-specific deletion of *Pparα* promotes NASH in the context of obesity"

### Supplemental Figure 4

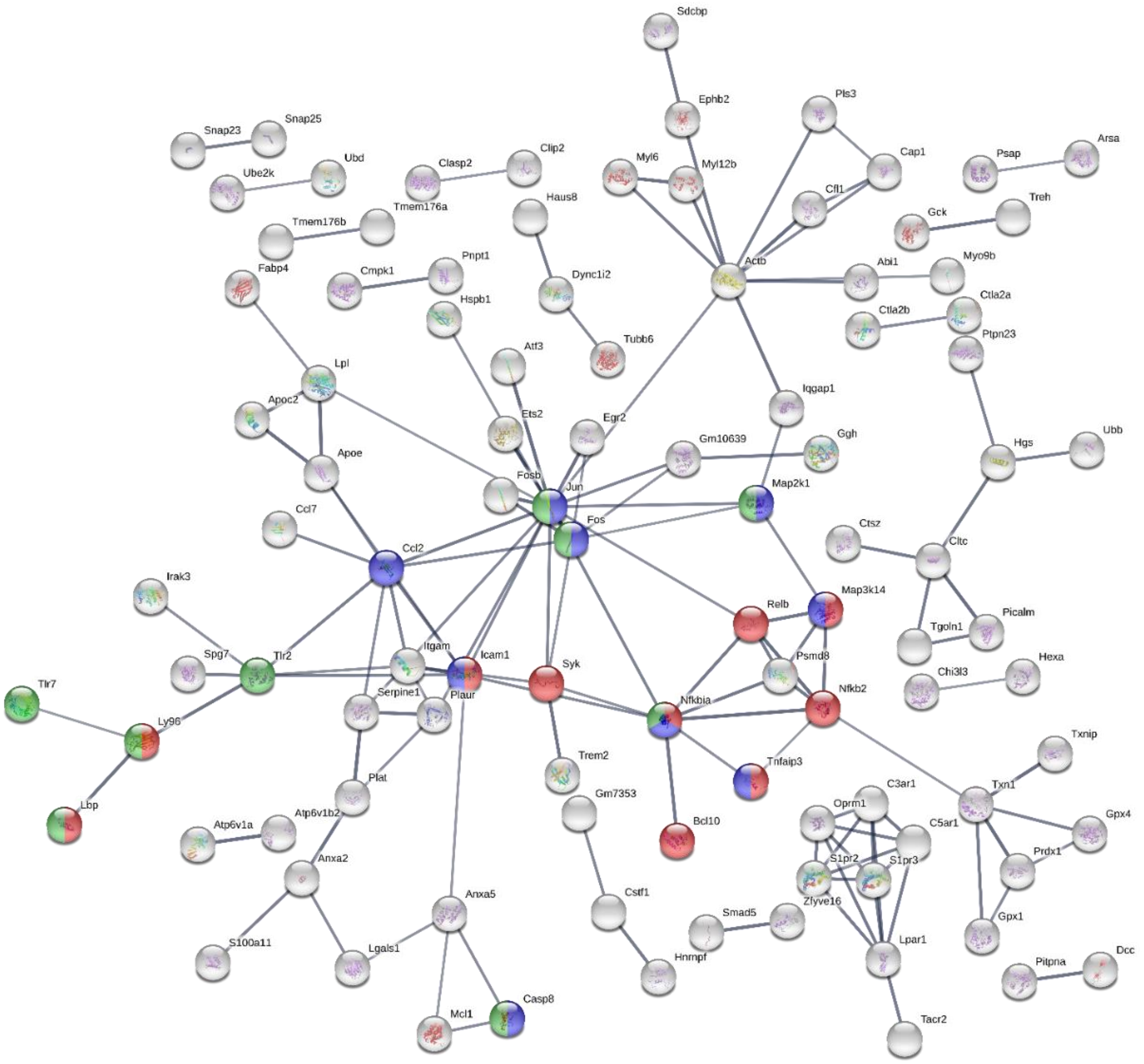

### Supplemental Figure 5

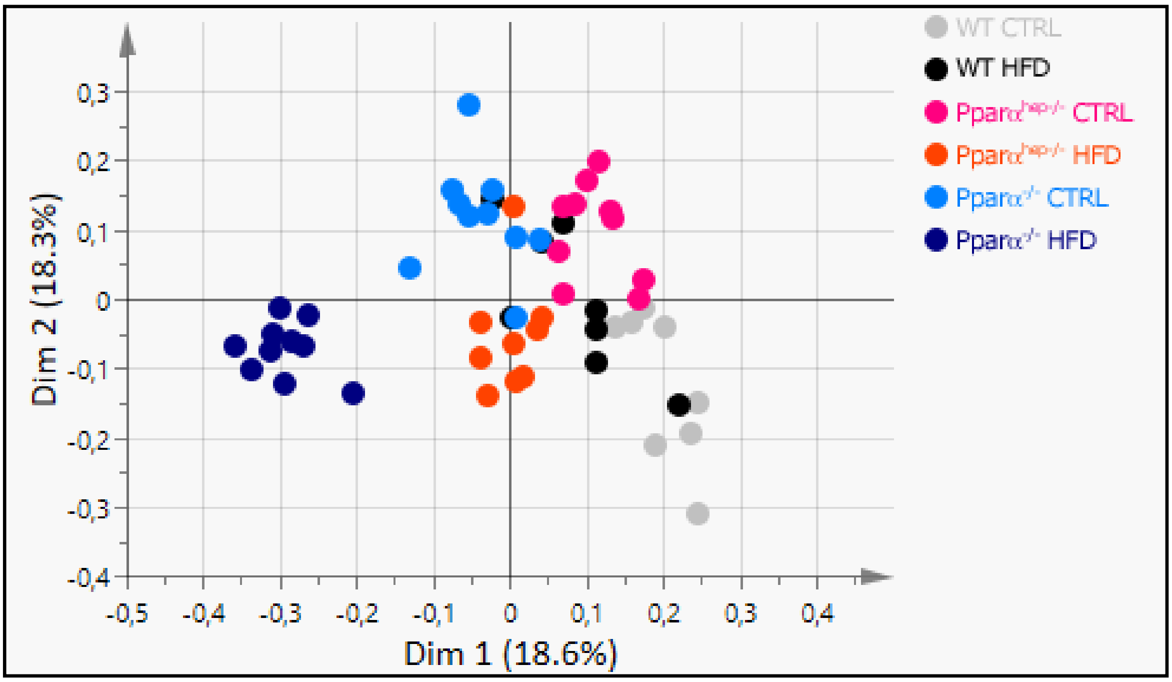
